# Predicting plant traits using large conglomerate RNA-seq datasets

**DOI:** 10.1101/2024.12.09.627626

**Authors:** John Anthony Hadish, Loren A. Honaas, Stephen Patrick Ficklin

## Abstract

RNA-seq datasets offer potential for predicting phenotypic traits, but the optimal dataset size for reliable predictions remains unclear. To explore this question, we compiled large-scale transcriptomic datasets across 12 plant species and used them to predict the phenotypic variables of tissue type and age. Predictions were made using random forest models. We created these models using an increasing number of samples to create performance curves.

These curves show that only a few hundred samples are required to achieve maximum accuracy for tissue classification, while predicting age demands a few thousand samples. Acceptable prediction accuracy can be achieved at even lower numbers of samples. Our findings provide a benchmark for designing transcriptomic studies aimed at phenotype prediction and highlight the differing complexities involved in predicting simple versus more complex traits.

**CORE IDEAS:**

- Plant transcriptomes are highly dynamic in response to internal and external conditions
- There is interest in using transcriptomes for predicting phenotypic traits and disorders
- It is unknown how many samples are required for accurate phenotype prediction
- We created 12 massive RNA-seq datasets and corresponding phenotypic data and used these to create prediction models
- Performance curves from our models provide insight into the required number of samples for accurate prediction

## 1 INTRODUCTION

DNA markers are used to associate genetic variation to a phenotype of interest. These have proven extremely useful for plant breeding and detecting risk factors in human health. In comparison to DNA, RNA is highly dynamic. Whereas DNA remains more or less unchanged over the course of an organism’s life, the quantity of RNA changes in response to both internal and external signals. It captures both the current and past states of an organism, reflecting its development and environmental experiences. This dynamic nature means that RNA offers potential for detecting and predicting phenotypes that are challenging to measure or exhibit delayed onset.

Transcriptomic (i.e., RNA) biomarkers are used in medical research with an emphasis on predicting cancer type and stage (Bostanci et al., 2023; Feng et al., 2019; Smith et al., 2020; Supplitt et al., 2021). Additionally, recent research has identified transcriptomic biomarkers in high-value agricultural crops, where researchers are interested in predicting traits such as flowering time (Azodi et al., 2020), flesh quality traits in apples, pears, and potatoes (Acharjee et al., 2016; Gapper et al., 2013; Hatoum et al., 2016; Leisso et al., 2016), and apple maturity (Favre et al., 2022). All of these traits and disorders are impacted by internal and external forces that cause RNA levels to change. These changes in RNA levels occur prior to the appearance of the trait or disorder.

Despite this growing interest in using transcriptomic biomarkers for predicting complex phenotypic traits, there has been little investigation into how many samples are required for accurate predictions. This lack of investigation is likely due to a lack of large transcriptomic datasets with corresponding phenotypes. To explore this question we created massive conglomerate transcriptomic datasets for 12 species of plants with corresponding phenotypic data for tissue type and age. We use these datasets to train random forest models. We created many models using increasing numbers of samples to establish performance curves for age and tissue. These performance curves can serve as a reference for selecting the number of transcriptomic samples required to accurately predict phenotypic traits.

## 2 MATERIALS AND METHODS

### 2.1 RNA-seq Data

All available RNA-seq data for 12 species of plants was retrieved from the National Center for Biotechnology Information (NCBI) Sequence Read Archive (SRA) (NCBI Resource Coordinators, 2016). Species were selected based on the quantity of data available and a desire to capture a wide diversity of plant species representing both model organisms and crop species. Species used for this project, in alphabetical order, were *Arabidopsis thaliana, Brachypodium distachyon, Glycine max, Gossypium hirsutum, Hordeum vulgare, Medicago truncatula, Oryza sativa, Populus trichocarpa, Solanum tuberosum, Triticum aestivum, Vitis vinifera* and *Zea mays*. Only RNA-seq data from Illumina sequencers (San Diego CA) was used.

RNA-seq samples were downloaded and gene expression quantified using a custom script: (https://gitlab.com/ficklin-lab/predicting-phenotypic-traits-using-conglomerate-rna-seq-datasets) which made use of fasterq-dump from the SRA toolkit (version 3.1.0) (NCBI SRA Toolkit Development Team., 2024) for sample retrieval. Kallisto (version 0.50.1) (Bray et al., 2016) was used for expression-level quantification. The workflow was performed on Washington State University’s High Performance Compute Cluster Kamiak. All reference transcriptomes were retrieved from JGI Phytozome (Goodstein et al., 2012) with genome version and references available in Supplemental Table 1. Primary transcripts for all genes were used. This processing resulted in a matrix where each row represents an RNA-seq experiment, each column represents the primary gene transcript, and the values represent the number of reads aligned to each gene transcript (i.e. gene expression). We refer to these files as gene expression matrices (GEMs).

After creation of a GEM for each species, filtering was performed. Genes with no recorded expression were removed, and samples with less than 1,000,000 reads were removed. After filtering, each GEM was normalized using mean ratio normalization (MRN) (Maza et al., 2013). We have made the raw GEMs available on Zenodo (DOI: 10.5281/zenodo.13328785).

### 2.2 Phenotypic Data

Phenotype annotations were retrieved from the NCBI BioProject database (Barrett et al., 2012; Federhen et al., 2014) using BioSampleParser (Limeta, 2020). Annotation terms were used inconsistently between different NCBI projects, so additional manual cleaning was required. For the tissue category, we classified samples as either “leaf”, “flower”, “root” or “other”. Common terms in the “other” category included derivatives of “shoot”, “seed”, and “seedling” but on further inspection we found that these terms were applied differently across projects and were therefore excluded.

We found that age was measured using a variety of different start points across NCBI projects (e.g. “days after flowering”, “days after sowing”, “days after inoculation”, etc.) and a variety of different units (e.g. hours, days, months). We changed all units to “days” and removed samples which were not measured from when the plant was sown.

In total, 69,138 samples for tissue, and 39,498 samples for age were manually classified into these categories for this project. A summary table by species of tissue type and age is available as Supplemental Table 2, with our classifications for each sample available on Zenodo (DOI: 10.5281/zenodo.13328785).

### 2.3 Random Forest Analysis

Random Forest Classification was used to evaluate the tissue dataset and Random Forest Regression was used to evaluate the age dataset (Breiman, 2001; Pedregosa et al., 2011). Our Python code is available here: (https://gitlab.com/ficklin-lab/predicting-phenotypic-traits-using-conglomerate-rna-seq-datasets). To summarize, both models were run with the parameters n_estimators=100, max_features=0.1, min_samples_leaf=3, bootstrap=True which were found to be high performing parameters from random search hyperparameter tuning. R^2^ was used to measure the performance of age (a continuous variable) and accuracy was used to measure the performance of tissue (a categorical variable). 100 repetitions were run for both tissue and age for each species. For each repetition, the data were first randomized with RNA-seq samples from the same bioproject always kept together (i.e., all in either the training dataset or all in the testing dataset) to prevent similar biological replicates being used to predict each other. After this randomization, models were created using increasing numbers of samples in the training dataset (e.g., 10, 30, 1000 samples etc.) and performance was recorded for each. After all replicates were run, the average performance was taken for each species at each of these sample sizes.

### 2.4 Best Fit Lines, Slope Calculations, and Percents

Best fit lines were fit to the random forest performance results using the R package nls2 (nls2 Development Team, 2024) with the Self Starting nLs logistic model (SSlogis)(R Core Team, 2022). We were interested mainly in identifying the point where performance increases leveled off so that we could compare the tissue and the age performance curves. We chose to fit these lines using only models which were run using at least 350 samples in the training dataset, as we found that this increased accuracy of the fit for this portion of the line. The derivative of these best fit lines was used to calculate the slope of the performance line. We chose the ad-hoc slope of 1e-5 to represent the point where no significant performance gains were seen in the model when adding more samples. We refer to this as the Point Of No Significant Performance Increase (PONSPI) in the remainder of the paper. We selected this slope to compare the performance increase of our two phenotypic treatments (tissue and age).

## 3 RESULTS

The average PONSPI for the tissue datasets was 662.98 samples (SD 238.54) and for the age dataset was 2248.90 samples (SD 55.37) (Figure 1, Supplemental Table 3). These two PONSPI means are statistically different (Welch’s t-test p = 8.26e-10).

**Figure 1.**
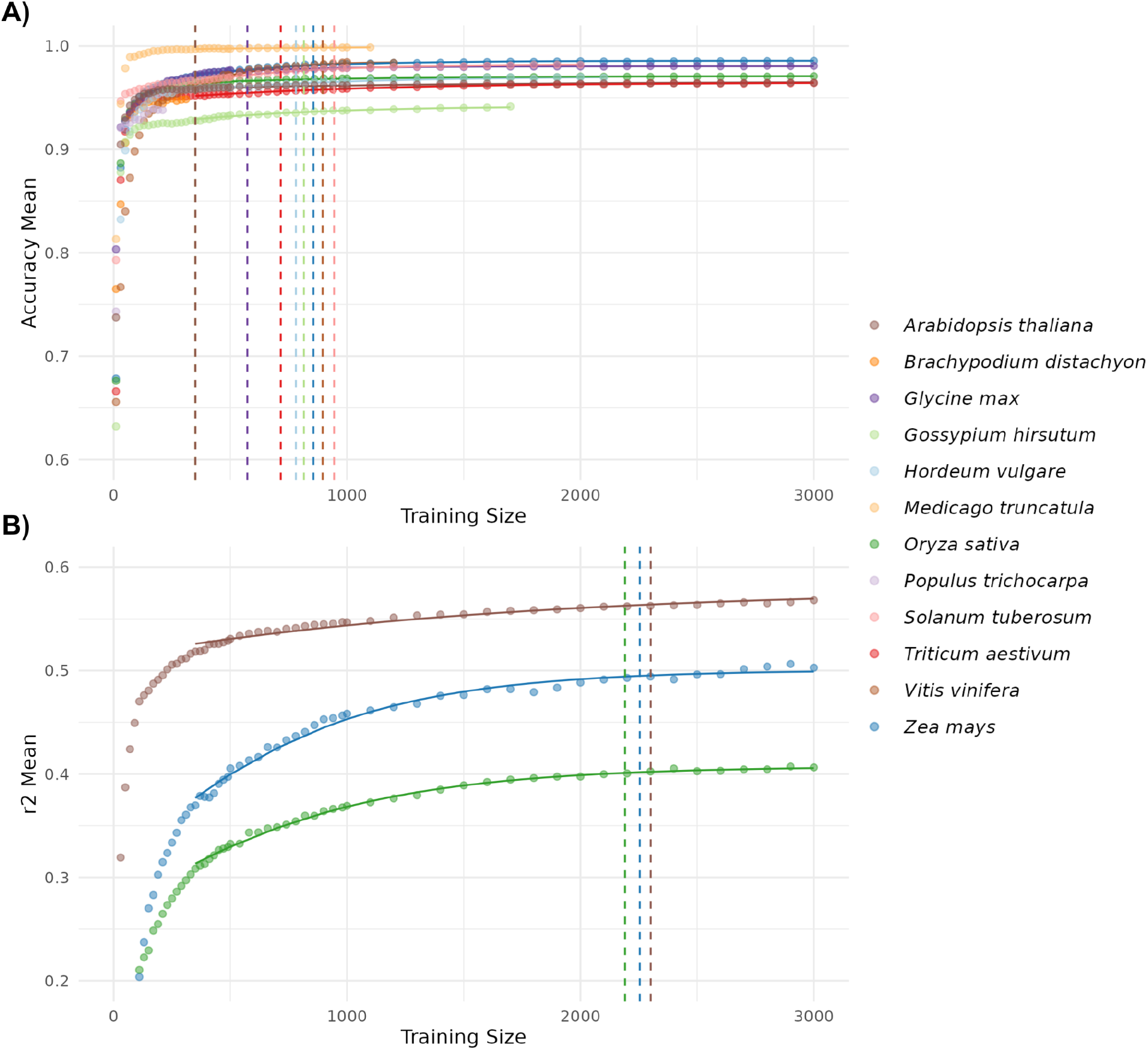
Model performance with increasing samples in the training set for **A)** Tissue and **B)** Age. The y-axis is performance measured by accuracy (for tissue) or R^2^ (for age). The x-axis represents the number of samples in the training dataset. Each point represents the average of 100 replicates. Vertical dashed lines represent the Point Of No Significant Performance Increase which we define as the derivative (slope) of the best fit line being under 1e-5. Performance values in table form for each species can be seen in Supplemental Table 3.

For the tissue dataset, 3 species, *Arabidopsis thaliana, Medicago truncatula*, and *Oryza sativa* reached PONSPI at the lowest possible point on the line of best fit (350 samples). Two species did not have enough samples to reach PONSPI (*Brachypodium distachyon* and *Populus trichocarpa*) (Supplemental Table 3). Performance results of the tissue classification were high, with the average peak performance on the testing dataset at 0.969 (SD 0.019) (See Supplemental Table 3 for performance by species).

For the age dataset, only 3 species gained an R^2^ score above 0.2 (*Arabidopsis thaliana, Oryza sativa*, and *Zea mays*) and were the only 3 used in the remainder of the analysis (Figure 1). Other species did not have performance that were considered better than random noise, with most receiving performance values below 0 R^2^ (Supplemental Table 3). The average R^2^ performance of the 3 species which met PONSPI was 0.499 (SD 0.088).

## 4 DISCUSSION

The goal of our research was to explore how many RNA-seq samples are needed to predict phenotypic outcome. This knowledge will help researchers interested in creating phenotypic prediction models using transcriptomic data. These models can then be used for identifying transcriptomic biomarkers for measuring difficult to measure or delayed phenotypic traits.

The two phenotypic traits we chose varied in their complexity, with the models created for tissue having only three categories (flower, leaf, and root) and the models created for age having a range of 11 to 74 unique timepoints. The tissue models performed well, likely due to the substantial transcriptomic differences between tissue types. In comparison, age was a more difficult phenotype to predict. This is likely partially due to differences in how age was recorded by researchers (it is obvious that a leaf if a leaf, but less obvious if age should be recorded as “days after sowing”, “days after germination”, “days after flowering”, etc.), and differences in experimental design (e.g. plants in lower light levels may be less physiological mature). It can also be partially explained in the more subtle differences (a plant which is 20 days old is similar to one which is 21 days old).

We would like to note that PONSPI was used to illustrate the differences in performance increase between our two phenotypic variables (age and tissue) and should not be thought of as the number of samples required for this type of analysis. The average PONSPI value shows that the performance of the tissue models increases rapidly with few samples whereas the performance of the age models increase at a more gradual rate. This allows us to see the difference in predicting two different complexity of traits. Very good results can be achieved with lower sample numbers and readers should refer to the performance curves in Figure 1 when deciding on the number of samples they will require for their model. Readers interested in creating their own predictive models should also consider how much transcriptomic variation is expected across their trait when selecting a sample size. More extreme transcriptome variation (such as in different tissue types) will allow for a lower number of samples compared to more subtle transcriptome variation (such as in age).

As the cost of a transcriptome sample continues to decrease it is feasible to see a future where thousands of samples could be regularly sequenced to develop machine learning models that can predict phenotypic outcome. Being able to associate gene expression in these samples with phenotypes allows for the monitoring of high value agricultural crops and the study of complex traits which are environmentally controlled. Our investigation of tissue and age demonstrated the feasibility of using transcriptomic data in random forest models, providing a foundation for predicting agriculturally important traits like abiotic and biotic stress. As more research is performed we will gain a more refined understanding of how many samples are needed.

We have made our datasets (both transcriptome and phenotypic annotations) for this experiment publicly available in the hope that they may prove useful for further analysis (https://zenodo.org/records/13328785).

## Abbreviations

PONSPI: Point Of No Significant Performance Increase

## ACKNOWLEDGMENTS

This research used resources of the Center for Institutional Research Computing at Washington State University

## SUPPLEMENTAL MATERIAL

**Supplemental Table 1:** Citation information on reference genomes used to map the RNA-seq data for each species.

**Supplemental Table 2:** RNA-seq sample counts for each species by phenotypic label.

**Supplemental Table 3:** Performance metrics of the testing dataset for each species for age and tissue prediction. The Point of No Significant Performance Increase (PONSPI) was set at the point where the derivative of the best fit line became less than 1e-5.

## DATA AVAILABILITY

DOI for Gene Expression Matrices and Phenotype Annotations: https://zenodo.org/records/13328785

Code: https://gitlab.com/ficklin-lab/predicting-phenotypic-traits-using-conglomerate-rna-seq-datasets

