## Supplemental Material for "Predicting plant traits using large conglomerate RNA-seq datasets"

**Supplemental Table 1:** Citation information on reference genomes used to map the RNA-seq data for each species.

| **Scientific Name** | Genome Version | Genome Reference |
|---|---|---|
| *Arabidopsis thaliana* | Tair10 annotations Araport 11 | (Cheng et al., 2017) |
| *Brachypodium distachyon* | v3.0 annotations v3.2 | (International Brachypodium Initiative, 2010) |
| *Glycine max* | Wm82.a6.v1 | (Schmutz et al., 2010) |
| *Gossypium hirsutum* | v3.1 | (Huang et al., 2020) |
| *Hordeum vulgare* | r1 | (Beier et al., 2017) |
| *Medicago truncatula* | Mt4.0v1 | (Tang et al., 2014) |
| *Oryza sativa* | version_7.0 | (Kawahara et al., 2013) |
| *Populus trichocarpa* | v4.1 | (Tuskan et al., 2006) |
| *Solanum tuberosum* | v6.1 | (Pham et al., 2020) |
| *Triticum aestivum* | IWGSC v2.1 | (Zhu et al., 2021) |
| *Vitis vinifera* | v2.1 | (Jaillon et al., 2007) |
| Zea mays | Zm-B73-REFERENCE-NAM-5.0.55 | (Jiao et al., 2017; Woodhouse et al., 2021) |

**Supplemental Table 2:** RNA-seq sample counts for each species by phenotypic label.

| **Scientific Name** | **Tissue Sample Count** | | | | **Age Sample Count** | |
|---|---|---|---|---|---|---|
| | **total** | flower | leaf | root | **total** | number of unique age values |
| *Arabidopsis thaliana* | **15316** | 2237 | 9195 | 3884 | **14546** | 42 |
| *Brachypodium distachyon* | **578** | 14 | 480 | 84 | **327** | 19 |
| *Glycine max* | **4847** | 267 | 2595 | 1985 | **1308** | 42 |
| *Gossypium hirsutum* | **2557** | 617 | 1252 | 688 | **672** | 22 |
| *Hordeum vulgare* | **4398** | 1818 | 1903 | 677 | **1569** | 25 |
| *Medicago truncatula* | **2026** | 1 | 426 | 1599 | **1299** | 27 |
| *Oryza sativa* | **11458** | 1991 | 6528 | 2939 | **5150** | 54 |
| *Populus trichocarpa* | **391** | 3 | 255 | 133 | **236** | 11 |
| *Solanum tuberosum* | **3421** | 64 | 1881 | 1476 | **1866** | 74 |
| *Triticum aestivum* | **8385** | 2515 | 4660 | 1210 | **4815** | 61 |
| *Vitis vinifera* | **3626** | 469 | 2896 | 261 | **1320** | 72 |
| *Zea mays* | **12135** | 1760 | 7078 | 3297 | **6390** | 111 |
| | 69138 | | | | 39498 | |

**Supplemental Table 3:** Performance metrics of the testing dataset for each species for age and tissue prediction. The Point of No Significant Performance Increase (PONSPI) was set at the point where the derivative of the best fit line became less than 1e-5.

| Species | Number of Samples Age PONSPI | Number of Samples for Tissue PONSPI | Max $R^2$ Age | Max Accuracy Tissue |
|---|---|---|---|---|
| *Arabidopsis thaliana* | 2301.251 | 350.000 | 0.582 | 0.966 |
| *Brachypodium distachyon* | | | -1.541 | 0.949 |
| *Glycine max* | | 572.823 | 0.046 | 0.981 |
| *Gossypium hirsutum* | | 814.865 | -0.291 | 0.942 |
| *Hordeum vulgare* | | 780.931 | -0.192 | 0.970 |
| *Medicago truncatula* | | 350.000 | -0.024 | 0.999 |
| *Oryza sativa* | 2190.941 | 350.000 | 0.407 | 0.971 |
| *Populus trichocarpa* | | | -0.517 | 0.939 |
| *Solanum tuberosum* | | 944.595 | -0.085 | 0.982 |
| *Triticum aestivum* | | 715.766 | 0.101 | 0.965 |
| *Vitis vinifera* | | 895.395 | -0.292 | 0.984 |
| *Zea mays* | 2254.505 | 855.405 | 0.506 | 0.986 |
